# A role for astrocytic miR-129-5p in Frontotemporal Dementia

**DOI:** 10.1101/2024.04.15.589528

**Authors:** Lalit Kaurani, Ranjit Pradhan, Sophie Schröder, Susanne Burkhardt, Anna-Lena Schuetz, Dennis M. Krüger, Tonatiuh Pena, Peter Heutink, Farahnaz Sananbenesi, Andre Fischer

## Abstract

Frontotemporal dementia is a debilitating neurodegenerative disorder characterized by frontal and temporal lobe degeneration, resulting in behavioral changes, language difficulties, and cognitive decline. In this study, smallRNA sequencing was conducted on postmortem brain tissues obtained from FTD patients with *GRN*, *MAPT*, or *C9ORF72* mutations, focusing on the frontal and temporal lobes. Our analysis identified miR-129-5p as consistently deregulated across all mutation conditions and brain regions. Functional investigations revealed a novel role of miR-129-5p in astrocytes, where its loss led to neuroinflammation and impaired neuronal support functions, including reduced glutamate uptake. Depletion of miR-129-5p in astrocytes resulted in the loss of neuronal spines and altered neuronal network activity. These findings highlight miR-129-5p as a potential therapeutic target in neurodegenerative diseases and also sheds light on the role of astrocytes in Frontotemporal dementia pathogenesis.

## Introduction

Frontotemporal dementia (FTD) is a devastating neurodegenerative disorder characterized by the progressive degeneration of the frontal and temporal lobes of the brain which leads to a wide range of symptoms, including changes in behavior, personality, and language, as well as cognitive decline [1] [2]. FTD affects individuals typically in the prime of their lives, striking those under the age of 65, making it one of the most common causes of early-onset dementia [3]. The clinical and genetic diversity of FTD presents a complex landscape [4]. While some cases occur sporadically, current data suggest that up to 48% of FTD cases have a familial basis [5]. Mutations in genes such as microtubule-associated protein tau (*MAPT*), progranulin (*GRN*), and chromosome 9 open reading frame 72 (*C9ORF72*) represent the most common genetic cause of FTD [5]. These genetic mutations are believed to disrupt critical cellular processes, eventually leading to neuronal cell death. Despite significant progress in understanding the genetic underpinnings of FTD, the development of effective therapies remains challenging [6]. In recent years, microRNAs (miRs) have emerged as important players in the molecular mechanisms underlying neurodegenerative diseases [7]. MiRs are 19-22 nucleotide-long RNA molecules that regulate protein homeostasis by binding to target mRNAs, leading either to their degradation or reduced translation [8]. Because one miR can affect a large number of mRNA targets that are often functionally linked, miRs have the ability to fine-tune gene expression and proteostasis and have been implicated in the regulation of multiple key pathways involved in brain function [9] [10] [11] [12]. Dysregulation of miRs has been observed in various neurodegenerative and neuropsychiatric diseases, including Alzheimer’s disease (AD), Parkinson’s disease (PD), amyotrophic lateral sclerosis (ALS), schizophrenia, and depression [13] [14] [15] [16] [17] [18] [19]. Today 2654 human miRs have been annotated [20] and microRNAome profiling in diseases gains increasing interest because changes in the levels of even one miR can indicate the presence of multiple pathologies [21] [22] [23]. Compared to neurodegenerative diseases such as AD, there is only limited data available on the role of miRs in FTD but some miRs have already been implicated in FTD pathogenesis. For example, miR-29b, miR-107, and miR-659 were implicated with *GNR* expression [24] [25] [26]. Additionally, miR-203 was detected as a putative hub microRNA controlling gene-expression networks de-regulated in models for Tau and Granulin pathology [27].

In our current study, we aimed to deepen the understanding of the role of miRs in FTD. Through comprehensive smallRNAome sequencing analysis of frontal and temporal cortex tissue samples obtained from FTD patients with mutations in *MAPT*, *GRN*, or *C9ORF72*, we uncovered a consistent down-regulation of miR-129-5p in all investigated brain tissues. Intriguingly, further investigations into the mechanistic implications of miR-129-5p loss, particularly in astrocytes, revealed its potential involvement in the disruption of neuronal plasticity, a hallmark of FTD.

Our study contributes to the growing body of knowledge on the intricate molecular landscape of FTD and highlights the critical role of miRs in this devastating disease. The data furthermore suggest that miR-129-5p could be a novel drug target to treat FTD and highlights the role of astrocyte dysfunction in disease progression.

## Materials and Methods

### Patient recruitment and clinical evaluation

Postmortem human brain tissue samples were acquired in accordance with a Material Transfer Agreement from the Netherlands Brain Bank and RNAseq analysis was approved by the ethical committee of the University Medical Center Göttingen (AZ 2/8/22 and AZ 29/9/18). Demographic information pertaining to the human brain specimens can be found in **Supplementary Table 1**.

### High-throughput sequencing of smallRNAomes

The NEBNext® small RNA library preparation kit was used to generate small RNAome libraries from 150 ng total RNA following the manufacturer’s guidelines. cDNA synthesis and PCR amplification were performed, and libraries were pooled. PAGE determined optimal size, and a 150-bp RNAome band was extracted for purification and quantification. Two nanomolar libraries were sequenced on the Illumina HiSeq 2000 platform using a 50-bp single-read setup. Demultiplexing utilized Illumina CASAVA 1.8, adapters were removed with cutadapt-1.8.1, and FastQC assessed sequence data quality.

### Processing and Quality Control

Reads were aligned to the Homo sapiens.GRCh38.p10 genome assembly (hg38) using the miRdeep2 package [28]}, with genome sequences accessed via the UCSC Genomic Browser. Bowtie-build tool (v1.12) mapped reads, and miRdeep2’s Perl-based scripts generated raw counts for microRNAs. Reads with fewer than 18 nucleotides were removed and used to quantify known microRNAs using miRDeep2’s quantifier.pl script.

### Differential Expression (DE) Analysis

Raw counts were used for DE analysis. Prior to DE analysis, raw read counts were log2-transformed and normalized for library size. Each sample was assigned a quality z-score, with samples of low quality (Z > 2.5 or Z < −2.5) considered outliers and excluded from further analysis. Read counts of 5 in at least 50% of the studied samples were used for subsequent DE analysis. The RUVSeq package [29] was employed to account for hidden batch effects and reduce unwanted variation. Data were corrected for age and gender. Differential expression analysis was performed using DESeq2, with microRNAs exhibiting a basemean >=50, and an adjusted p-value of < 0.05 considered differentially expressed.

### Gene ontology enrichment and pathway analysis

The Gene Regulatory Network (GRN) for miRNA-target genes was built using validated targets from miRTarBase (v7.0). Target genes with brain expression were selected using GTEx portal (https://gtexportal.org/home/). Genes demonstrating a moderate level of expression were prioritized. Gene expression in the GTEx portal is quantified using Transcripts Per Million (TPM), a reliable metric for identifying genes with moderate expression levels. A cutoff of 10 TPM was employed to determine moderate to high level expression in the brain. As such, the final list of target genes selected for pathway analysis was not only potentially targeted by candidate miRNAs, but also demonstrated a meaningful level of expression in the brain according to GTEx portal data. This approach of dual-level filtering ensured that subsequent pathway analysis was based on biologically relevant and expressed genes. Then Cytoscape 3.2.1’s ClueGO v2.2.5 plugin was used to determined biological processes and pathways. Significance was calculated using a two-sided hypergeometric test and Benjamini-Hochberg adjustment. KEGG and Reactome databases informed pathway analysis, and GRN was generated for deregulated mRNAs. Processes and pathways with an adjusted p-value of < 0.05 were further evaluated. Later using the systematic approach, the key biological processes were selected primarily based on their Gene Ontology (GO) levels. This GO level is a systematic method used to describe the attributes of genes and gene products, such as cellular components, molecular functions, and biological processes. For instance, a lower GO level signifies a more specific function of genes, while a higher GO level embodies biological processes with more generalized functions. Biological processes were ranked according to their GO levels in a hierarchical manner, with lower GO levels (indicating more specific gene function) being given higher priority. Following this categorization, biological processes with an adjusted p-value of less than 0.05 were selected to ensure statistical significance and to mitigate the possibility of false positives. Certain cancer-related pathways if appeared in top 10 significant processes; however, they were not incorporated into our further analysis. Given our primary focus on Frontotemporal Dementia (FTD), a neurodegenerative disease, we consciously directed our efforts towards the exploration of target genes that are implicated in FTD pathogenesis. As such, only those biological processes and pathways that bore relevance to neurodegenerative diseases, with a particular emphasis on FTD, were retained for in-depth examination.

The network shonw in Fig S2 was built using Cytoscape (v3.7.2) based on automatically created lists of pairwise interactors. We used inhouse Python scripts to screen interactome databases for annotated interactions using miRNA-129-5p and the list of up- and down-regulated genes as input. Results thus obtained contain interactions between input and all (non-)coding genes as annotated in the databases. Interaction information was collected from six different databases: NPInter, RegNetwork, Rise, STRING, TarBase, and TransmiR. The lists of pairwise interactors were loaded into Cytoscape to built a network which was visually modified according to up- and down-regulated genes. The initial network was truncated to a core network with PathLinker (v1.4.3) using 5000 paths and the input list as source.

### Quantitative PCR

The miScript II RT Kit synthesized cDNA from 200 ng total RNA, which was used for qPCR analysis of both mRNA and microRNA. MicroRNA-specific forward and universal reverse primers were used, with U6 small nuclear RNA as a control. Gene-specific primers quantified mRNA, normalized against GAPDH. The 2–^ΔΔCt^ technique calculated fold changes, and a Light Cycler® 480 Real-Time PCR System performed qPCR. Primers sequences are provided in the **supplemental table 12**.

### Preparation of microRNA lipid nanoparticles

miR-129-5p expression was inhibited using miR-129-5p inhibitor sequences (anti-miR-129), an antisense oligos (ASO), and negative control sequences (scramble control) from Qiagen. The Neuro9^TM^ siRNA SparkTM Kit produced lipid nanoparticle (LNP) formulations for microRNA inhibitors or ASOs (5 nmol) using a proprietary lipid blend. NanoAssemblr^TM^ Spark^TM^ technology encapsulated miRNA inhibitors or ASOs in a microfluidic device with controlled mixing conditions (Precision Nanosystems). Following the manufacturer’s instructions, 5 nmol lyophilized microRNA inhibitors or ASOs were reconstituted, diluted, and encapsulated using the NanoAssembler Spark system.

### Primary neuronal culture

Primary neuronal cultures were prepared from E17 CD1 mice (Janvier Labs, France). Mice were sacrificed, and embryos’ brains, meninges, and cortices and hippocampi were dissected and rinsed with PBS. Following trypsin and DNase incubation, single-cell suspensions were obtained. Cells were plated on poly-D-lysine-coated 24-well plates with Neurobasal media and B-27 supplement. Primary hippocampal neurons were used in experiments at DIV10-12.

### Dendritic spine labeling

Primary hippocampal neurons and neuron-astrocyte co-cultures were fixed with 2% PFA and dendritic spines were labeled using Dil dye (Life Technologies-Molecular Probes). After a 10-minute incubation and thorough rinsing, cells were incubated overnight, cleaned, and mounted with mowiol. High-magnification images were captured using a multicolor confocal STED microscope with a 60x oil objective. Spine density and total spine length were measured with ImageJ.

### Multi-electrode assay

E17 embryos were used for neuronal culture in neurobasal media (Gibco, USA). After centrifugation, cells were suspended, combined with laminin (Merck, Germany), and plated in MEA plates with 16 electrodes per well, pre-coated with PDL. Media was changed every third day, and at DIV10, basal spontaneous activity was measured. Cells were treated with LNPs containing miR-129-5p inhibitors (anti-miR-129) or scramble sequence as controls. Spontaneous neural activity was monitored from DIV12 using the Maestro Apex Platform (Axion Biosystems, USA). Data was recorded for 29 hours, retrieved, and analyzed with AxlS Navigator (Axion Biosystems, USA). Values were plotted and analyzed using Prism 8.3.1 (GraphPad Software LLC, United States). The calculation of the Neural Activity Score (NAS) was performed to quantify the overall neural activity from the array of neural metrics acquired. This calculation was conducted using a Principal Component Analysis (PCA) approach implemented with the ‘pca’ function from the ‘pcaMethods’ package in R. This method enables the reduction of the complex high-dimensional data into fewer dimensions, thereby encapsulating the majority of the data’s variance in a more manageable form. Our PCA model was built on the z-score normalized neural metrics. This model enabled us to derive a set of orthogonal (i.e., uncorrelated) principal components that maximally explain the variance within the neural metrics. The NAS was then calculated as the first principal component of this PCA model, capturing the largest possible variance from the neural metrics. The NAS thus represents a composite measure of neural activity based on the largest common pattern of variation within the measured neural metrics.

### Primary astrocyte culture

P0 to P2 mice pups of CD1 background were sacrificed by decapitation and their cortex dissected, careful to remove all meninges from each hemisphere. 2 brains were pooled per 15-mL falcon containing ice-cold PBS. Tissue was dissociated in a pre-warmed solution of 0.05% trypsin-EDTA (in PBS) at 37°C for 20-minutes. Trypsinization was stopped by adding a processing medium (10 mM HEPES in HBSS) and cells were washed twice to wash off remaining trypsin-EDTA. Dissociated tissues were homogenized in the processing media, filtered through a 100μm cell strainer, and transferred into a coated T75 flask. The day of plating the cells was considered as day 0 (DIV0). On the next day, media was completely exchanged for a new processing medium and three days later half the medium was changed. At DIV 7, astrocytes were placed on a shaker at 110rpm, at 37°C, for 6-hours. After, cells were washed with preheated PBS, followed by an incubation of 5-minutes with 0.25% trypsin-EDTA. Trypsinization was once again stopped by adding processing medium and cells were loosened by forceful pipetting. Cell suspension was then centrifuged at 3,200 g for 4-minutes at 20°C. Cell pellet was resuspended in 1mL MM+ media (Neurobasal Plus Medium, 2% B27, 1% P/S, 1% GlutaMAX) containing 5ng/mL HBEGF (Sigma-Aldrich, REF. 4643). Cells were counted in a Neubauer counting chamber and seeded at a density of 60,000 cells/well in 12-well plates that had been previously coated with 0.05 mg/L poly-D-Lysine. Half the cell medium was exchanged for fresh MM+/HBEGF media once a week.

### Glutamate uptake assay

Primary astrocytes were grown in 96 well plates and treated for 48 hours. On DIV14, the medium was removed and the cells were washed with HBSS for 10 min at 37°C. The HBSS was removed and the cells were incubated with 100 µM glutamate in HBSS for 1 hour at 37°C. Then, the supernatant was collected and the amount of glutamate in the supernatant was determined using the Glutamate Glo Assay (Promega).

### Co-culture system

A confluent layer of neurons was established in multi-wells as previously described. Astrocytes were cultured in a T75 flask and transfected with either anti-miRNA-129-5p or scramble control at DIV 10. At neuronal DIV 12 and astrocytic DIV 12, LNP-treated astrocytes were split and seeded onto neurons at a density of 20% (60,000 astrocytes/well) of neuronal density. Co-cultures were maintained for an additional 48h, then used for RNA sequencing.

### RNA sequencing

Total RNA was employed for library preparation using the TruSeq RNA Library Prep Kit v2 (Illumina, USA), following the manufacturer’s instructions. A starting material of 500 ng RNA was used. Library quality was assessed using a Bioanalyzer (Agilent Technologies), while library concentration was determined with the Qubit™ dsDNA HS Assay Kit (Thermo Fisher Scientific, USA). Multiplexed libraries were directly loaded onto a HiSeq2000 (Illumina) with a 50 bp single-read configuration. Demultiplexing was performed using Illumina CASAVA 1.8, and sequencing adapters were removed utilizing cutadapt-1.8.1.

### Publicly available datasets

This study uses various publicly available datasets to investigate the cell type-specific expression of candidate microRNAs and differentially expressed genes. Gene expression specific to neurons, astrocytes, and microglia was investigated using published single-cell data [30]. miR-129-5p and miR-212-5p expression in human neurons and astrocytes was investigated using published cell type specific data [31].

### Statistical analysis

Statistical analysis was conducted using GraphPad Prism version 8.0. Data are presented as mean ± standard deviation or as boxplots, with each ‘n’ representing a biological sample. Analyses were performed employing a two-tailed unpaired t-test. Cell-type-specific gene overlap analysis was conducted using the GeneOverlap package in RStudio (v1.4.1106). Enriched gene ontology and pathway analyses were executed using Fisher’s exact test and Benjamini-Hochberg correction for multiple comparisons.

## Results

### Altered microRNA expression in postmortem tissue samples of FTD patients

We conducted small RNA sequencing on postmortem brain tissue obtained from two distinct brain regions – the frontal and temporal lobes – of patients with mutations in the MAPT (n = 13), GRN (n = 6), or C9ORF72 (n = 8) genes causing FTD. Additionally, brain tissue from non-demented controls (n = 16) was included in the analysis **(Fig 1A; Supplemental Table 1)**. For three individuals, we lacked frontal lobe samples, resulting in a total of 77 small RNA sequencing datasets. Using differential expression analysis (with adjusted p-value < 0.1, basemean >= 50, log2FC = +/− 0.2), we identified significant changes in miRNA expression between control and FTD cases in both analyzed brain regions **(Fig 1B, C; Supplemental Table 2)**. Notably, a greater number of miRNAs showed alterations in the frontal lobe (n = 276) compared to the temporal lobe (n = 130; **Fig 1D**). Further analysis revealed that the number of deregulated microRNAs was similar among *C9ORF72*, *GRN*, and *MAPT* carriers in the frontal lobe, whereas only a few microRNAs exhibited significant alterations when comparing C9ORF72 or GRN carriers to controls in the temporal lobe **(Fig 1E)**. Since disease-associated changes in microRNA levels are believed to reflect a response to altered proteostasis and mRNA levels, these findings could suggest that the observed variations across brain regions and patient groups may be attributed to different patterns in the occurrence of pathology as it has been observed in FTD cases [32].

**Fig. 1.**
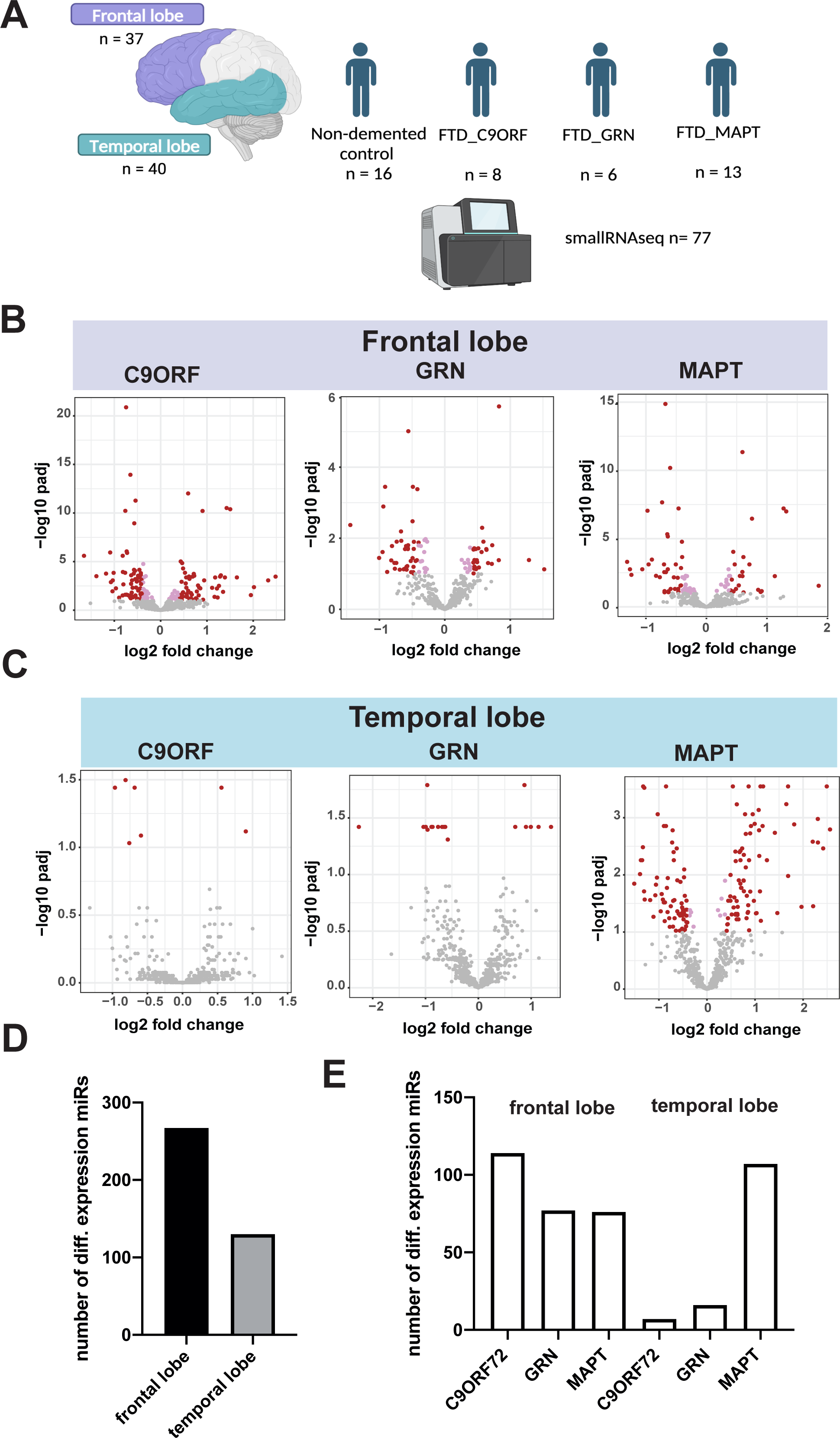
Differential microRNA expression the frontal and temporal lobe of FTD patients. **A.** Schematic representation of the study design**. B**. Volcano plot showing microRNA expression in the frontal lobe of control vs FTD patients with *C9ORF72*, *GRN* or *MAPT* mutations. The plot illustrates the relationship between log2 fold change (log2FC) and statistical significance (-log10 of the adjusted *p*-value, padj). **C.** Volcano plot showing microRNA expression in the temporal lobe of control vs FTD patients with *C9ORF72*, *GRN* or *MAPT* mutations. The plot illustrates the relationship between log2 fold change (log2FC) and statistical significance (-log10 of the adjusted *p*-value, padj). **D.** Bar plot illustrating the total number of differentially expressed microRNAs in the frontal and temporal lobes, obtained by summing the data from all three mutations. **E.** Bar plot depicting the number of differentially expressed microRNAs in the frontal and temporal lobes for each individual FTD mutation.

Since all three FTD mutations eventually lead to similar traits in patients, our main goal was to identify commonly disrupted miRNAs in both the frontal and temporal lobes across all patient groups. Initially, we compared miRNAs with significant alterations in the frontal lobe of patients carrying the *C9ORF72*, *GRN*, or *MAPT* mutation, resulting in the discovery of 30 commonly disrupted microRNAs **(Fig. 2A, supplemental table 3)**. Similarly, we found 5 deregulated miRNAs in all conditions in the temporal lobe **(Fig. 2B, supplemental table 3)**.

**Figure 2.**
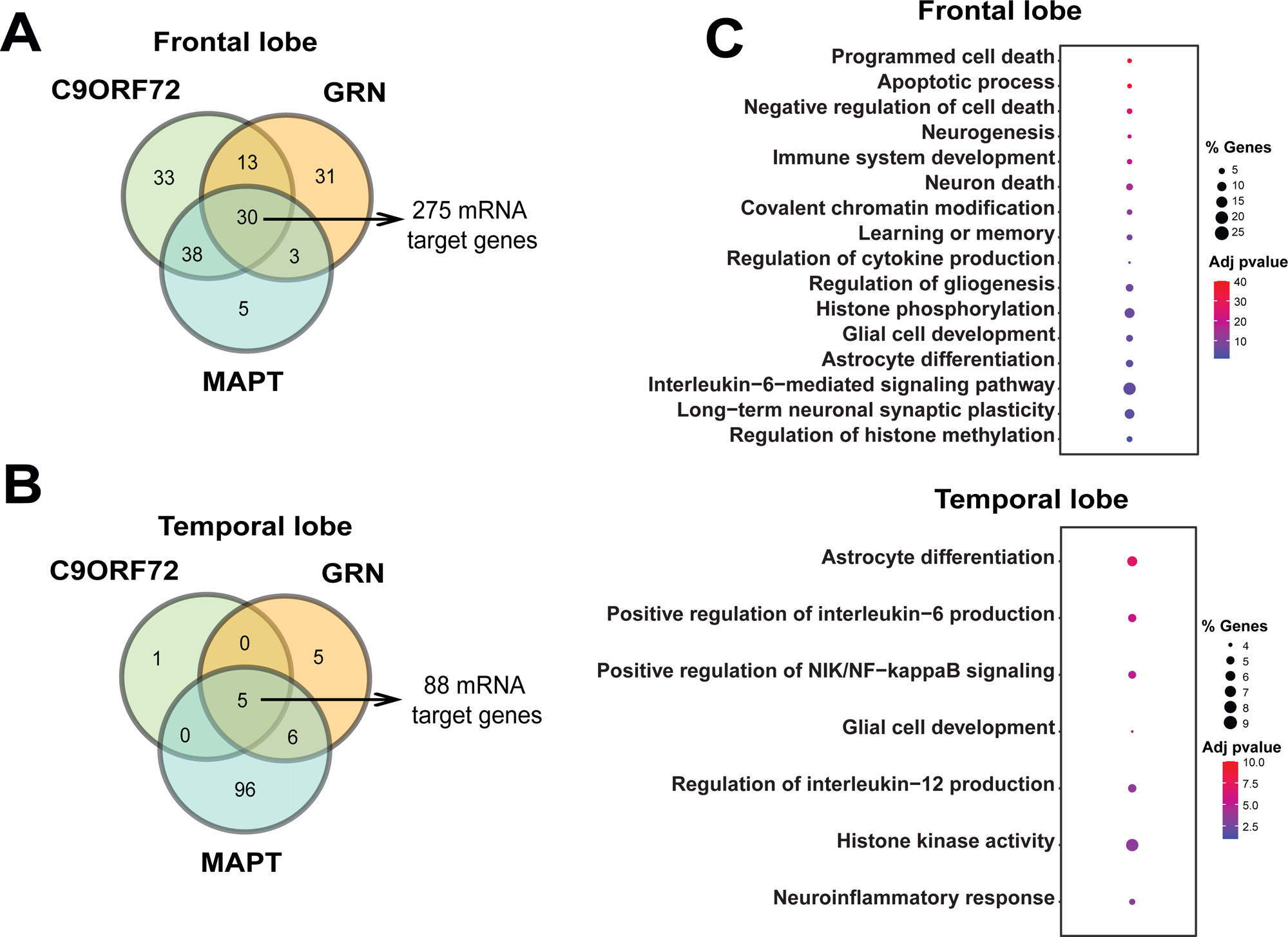
GO terms for the target genes of microRNAs commonly deregulated in the frontal and temporal lobe of FTD patients. **A.** Venn diagram showing that 30 microRNAs (see also supplemental table 3) are commonly deregulated in the frontal lobe of all analyzed samples. **B.** Venn diagram showing that 5 microRNAs (see supplemental table 3) are commonly deregulated in the temporal lobe of all analyzed samples. **C.** Representative GO terms based on the commonly deregulated miRNAs in the frontal lobe (upper panel) and the temporal lobe (lower panel).

We utilized the miRTarBase database to identify mRNA targets of the commonly disrupted miRNAs in both lobes. Subsequently, we narrowed down the target mRNAs to those expressed in the human brain using the GTEx portal, which led to the identification of 275 potential target mRNAs **(supplemental table 4**).

Analysis of Gene Ontology (GO) terms associated with these mRNAs revealed several pathways related to neuronal cell death, such as “programmed cell death,” “apoptotic processes,” “negative regulation of cell death,” and “neuron death” **(Fig. 2C)**. Additionally, significant processes were linked to inflammatory responses (e.g., “immune system development,” “negative regulation of immune system process,” “regulation of cytokine production,” “interleukin-6-mediated signaling pathway”), synaptic plasticity, and memory function (e.g., “neurogenesis,” “learning or memory,” “regulation of long-term neuronal synaptic plasticity”). Furthermore, processes associated with glial cells, particularly astrocytes, were identified (e.g., “regulation of gliogenesis,” “glial cell development,” “astrocyte differentiation”), along with processes related to chromatin regulation (e.g., “covalent chromatin modification,” “histone phosphorylation,” “regulation of histone methylation”). These findings suggest that the deregulated miRs are closely associated with neurodegenerative processes observed in FTD patients and relevant model systems [33]. The complete list of GO terms and their corresponding genes can be found in supplemental table 5 **(supplemental table 5)**.

We applied the same approach to gain a deeper understanding of the potential role of the five commonly deregulated miRs in the temporal lobe of FTD patients. We identified 88 miR targets that are enriched in the brain **(supplemental table 4)**. Similar to the findings in the frontal lobe data, GO term analysis of these targets revealed processes associated with neuroinflammation, such as “positive regulation of interleukin-6 production,” “positive regulation of NIK/NF-kappaB signaling,” “regulation of interleukin-12 production,” and “neuroinflammatory response.” Additionally, processes related to glial cells, including astrocytes (“astrocyte differentiation”, “glial cell development”), and chromatin regulation (“histone phosphorylation”, “histone kinase activity”) were identified (**Fig. 2C**, **also see supplemental table 5**).

Interestingly, five identical GO terms were detected in both the frontal and temporal lobe datasets, namely “regulation of interleukin-12 production,” “glial cell development,” “astrocyte differentiation”, “cellular response to amyloid-beta,” and “histone phosphorylation.” The analysis of the temporal lobe data did not yield significant GO terms related to neuronal cell death, even when considering all 144 significant GO terms identified **(supplemental table 5)**. This may reflect a difference in the onset of neuronal cell death between the frontal and temporal lobes of FTD patients, consistent with previous literature [32]. In this scenario, the changes observed in the temporal lobe may reflect an early stage of pathology. It is interesting to note that the most significant GO terms were linked to astrocyte function and neuroinflammation **(see Fig. 2C)**.

In addition to the miRs commonly regulated in the frontal and temporal lobes across patient groups, we also analyzed the miRs specific to each condition using the same approach as described above (**supplemental table 6)**. While the data revealed that distinct GO terms were enriched for the specific conditions, it is worth highlighting that “miRNA processing” was amongst the top 10 GO terms affected in the frontal lobe of patients with MAPT mutations, **(Fig. S1, supplemental table 7)**.

Two microRNAs, miR-129-5p and miR-212-5p, exhibited consistent deregulation across all patient samples, regardless of the specific mutation or brain region **(Fig. 3A)**. We validated the decreased expression of these microRNAs via qPCR **(Fig. 3 B,C)**. Extensive research has focused on miR-212-5p in the adult brain. Reductions in its levels have been associated with memory impairment in mice [34] [35] [36], and miR-212-5p is known to be decreased in several neurodegenerative diseases such as Alzheimer’s disease (AD), Huntington’s disease, and Parkinson’s disease (PD) [37] [38] [39] [40] [41] [42] [43] [44] [45]. Additional studies suggest that targeting miR-212-5p expression is neuroprotective in models of PD, AD, and traumatic brain injury [46] [47] [48]. MiR-212-5p forms a cluster with miR-132 [36], which has also been linked to neurodegenerative diseases. MiR-132 has been found to be decreased in the brains of AD and PD patients [49] [16] as well as in FTD patients [50]. In our dataset, miR-132-3p was commonly down-regulated in the temporal region of C9ORF, GRN, and MAPT mutant carriers, while in the frontal region, miR-132-3p was only decreased in patients with C9ORF mutations **(supplemental table 2)**.

**Figure 3.**
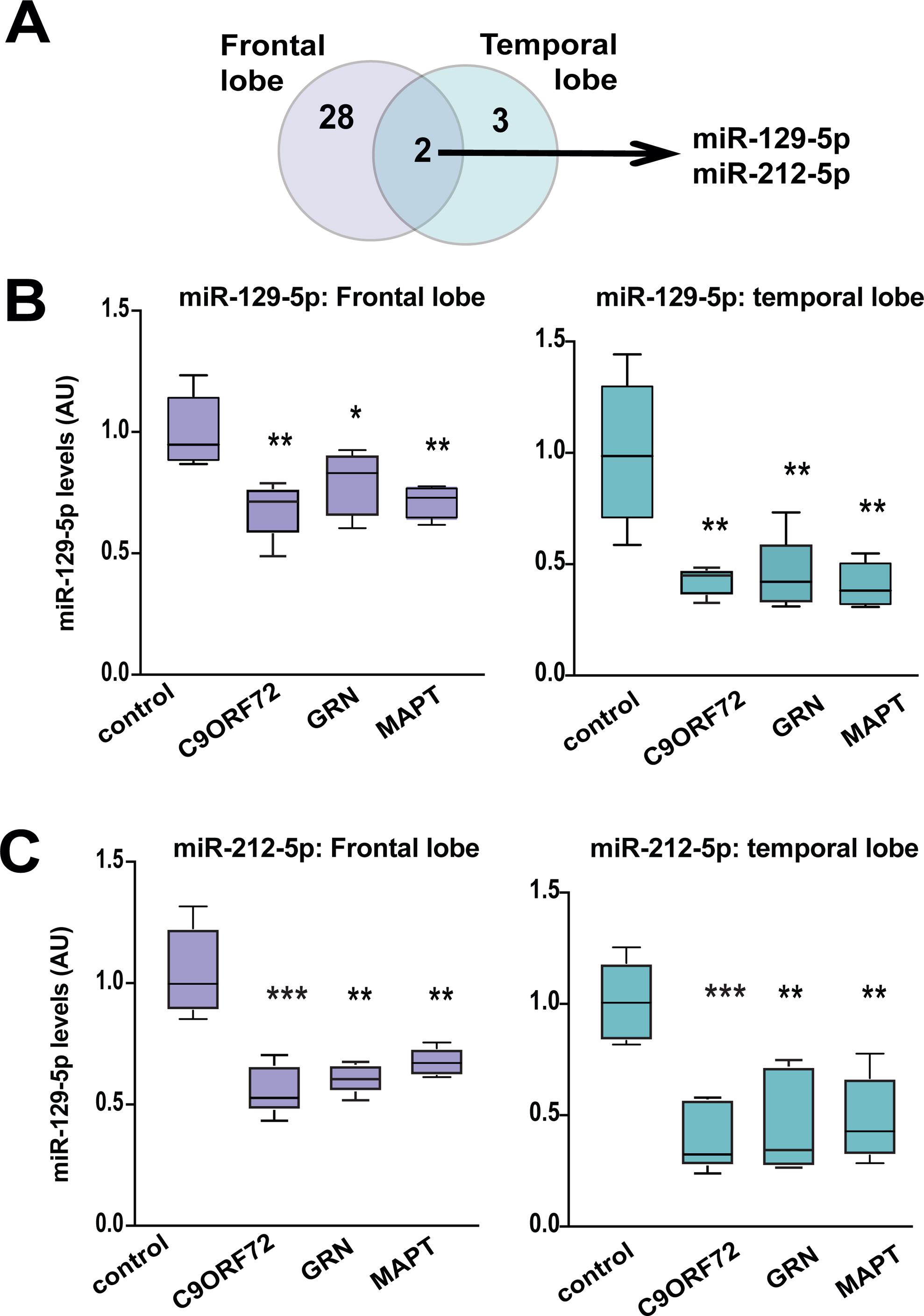
Deregulation of miR-212-5p and miR-129-5p in FTD. **A.** Venn diagram showing that miR-212-5p and miR129-5p are commonly deregulated in FTD patients. **B.** The qPCR results depict the expression of miR-129-5p in the frontal lobe (Left panel) and temporal lobe (Right panel) of both control and FTD patients (n=5/group). **C.** The qPCR results depict the expression of miR-212-5p in the frontal lobe (Left panel) and temporal lobe (Right panel) of both control and FTD patients (n=5/group). The horizontal line in the box plot represents the median, the box spans 25 and 75% quantile, and the whiskers represent the smallest and largest values in the 1.5x interquartile range. (*P < 0.05; **P < 0.01, ***P < 0.001; *t*-test unpaired; 2-tailed)

In summary, these data indicate that decreased levels of the miR132/212 cluster are common across several neurodegenerative diseases and are associated with impaired cognitive function.

In comparison to miR-212-5p, there is limited knowledge regarding the role of miR-129-5p in neurodegenerative diseases. A recent study observed decreased miR-129-5p levels in postmortem brains samples of AD patients from the ROS/MAP cohort and a negative correlation to cognitive function [51], thereby confirming previous findings [17] [52]. Another study reported decreased levels of miR-129-5p in postmortem brains from PD patients [16]. To the best of our knowledge, altered miR-129-5p levels have not been reported in FTD. At the functional level miR-129-5p has been linked to epileptic plasticity and homeostatic synaptic downscaling in excitatory neurons [53].

In summary, existing literature strongly supports the down-regulation of miR-212-5p and miR-129-5p in neurodegenerative diseases, including Alzheimer’s disease (AD) and Parkinson’s disease (PD). Our study adds to this body of evidence by demonstrating a common down-regulation of miR-212-5p and miR-129-5p in two different brain regions of FTD patients carrying either the *C9ORF72*, *GRN*, or *MAPT* mutations. These findings suggest that both microRNAs control key cellular pathways central to various neurodegenerative diseases. Therefore, conducting functional analysis on these microRNAs becomes an important research topic. Such analysis could provide valuable insights to guide the development of novel therapeutic strategies for treating these diseases.

While miR-212-5p’s role in the adult brain is well-studied, there is comparatively limited understanding of the function of miR-129-5p in this context. Therefore, we have chosen to investigate this specific microRNA further within the scope of our current study

### Knockdown of miR-129-5p affect neuronal plasticity and induces a gene-expression signature linked to neuroinflammatory processes

We utilized existing microRNA expression datasets [54] to analyze miR-129-5p levels in neurons, microglia, and astrocytes. The data revealed high expression of miR-129-5p in neurons and astrocytes compared to microglia (refer to **Fig. 4A**). These findings are consistent with the Gene Ontology (GO) terms associated with altered miRNA expression in our datasets, where pathways such as “astrocyte differentiation” were among the top-ranking, alongside pathways linked to “neuron death” (see **Fig. 2**). To gain further insight into the role of miR-129-5p, we transfected mouse primary neuronal mixed cultures (PNM cultures), which contain mainly neurons and astrocytes, with antisense miR-129-5p locked nucleic acid (anti-miR-129). As a control, we used a scrambled sequence (sc-control). The anti-miR-129 construct significantly reduced the detection of miR-129-5p levels in PNM cultures when assayed 48 hours after transfection **(Fig. 4B)**. Next, we analyzed the impact of anti-miR-129 administration on synaptic morphology in PNM cultures. The number of dendritic spines was significantly decreased in PNM cultures treated with anti-miR-129 compared to the sc-control group **(Fig. 4C)**. To investigate if the altered morphology affects neuronal network plasticity, we performed a multielectrode array (MEA) assay on PNM cultures. Basal activity was measured at DIV 10, followed by treatment with anti-miR-129-5p or sc-control oligonucleotides for 48 hours before transferring the plates to the MEA device for further recording. Anti-miR-129-5p treated PNM cultures exhibited significantly reduced neuronal activity compared to the sc-control group, including parameters such as the weighted mean firing rate and the number of network bursts **(Fig. 4D)**. To elucidate the molecular processes controlled by miR-129-5p further, we performed RNA sequencing of PNM cultures upon miR-129-5p knockdown. PNM cultures treated with sc-control oligonucleotides were used for comparison. Analysis of gene expression revealed 333 deregulated genes (adjusted p-value <0.05, log2foldchange = ±0.4, basemean >= 50), comprising 155 upregulated and 188 downregulated genes (refer to **Fig. 4E**, **Supplementary table 8**). Differential expression of selected genes was confirmed via qPCR **(Fig. 4F)**. GO term analysis of upregulated genes indicated overrepresentation of key biological processes linked to inflammation, such as ‘response to interferon-gamma’, ‘glial cell proliferation’, including ‘astrocyte development’, or ‘positive regulation of phagocytosis’ **(Fig. 4G)**. GO term analysis of downregulated genes revealed several significant pathways linked to ‘DNA replication’, ‘regulation of oligodendrocyte differentiation’, ‘neuron fate commitment’, ‘regulation of Notch signaling pathway’, ‘transmembrane transporter activity’, or ‘astrocyte differentiation’ **(Fig. 4G; Supplementary Table 9)**. Since PNM cultures consist mainly of neurons and astrocytes, we analyzed the differentially expressed genes for their cell-type expression pattern and noticed that the deregulated genes were mainly associated with astrocytes **(Fig. 4H)**.

**Figure 4:**
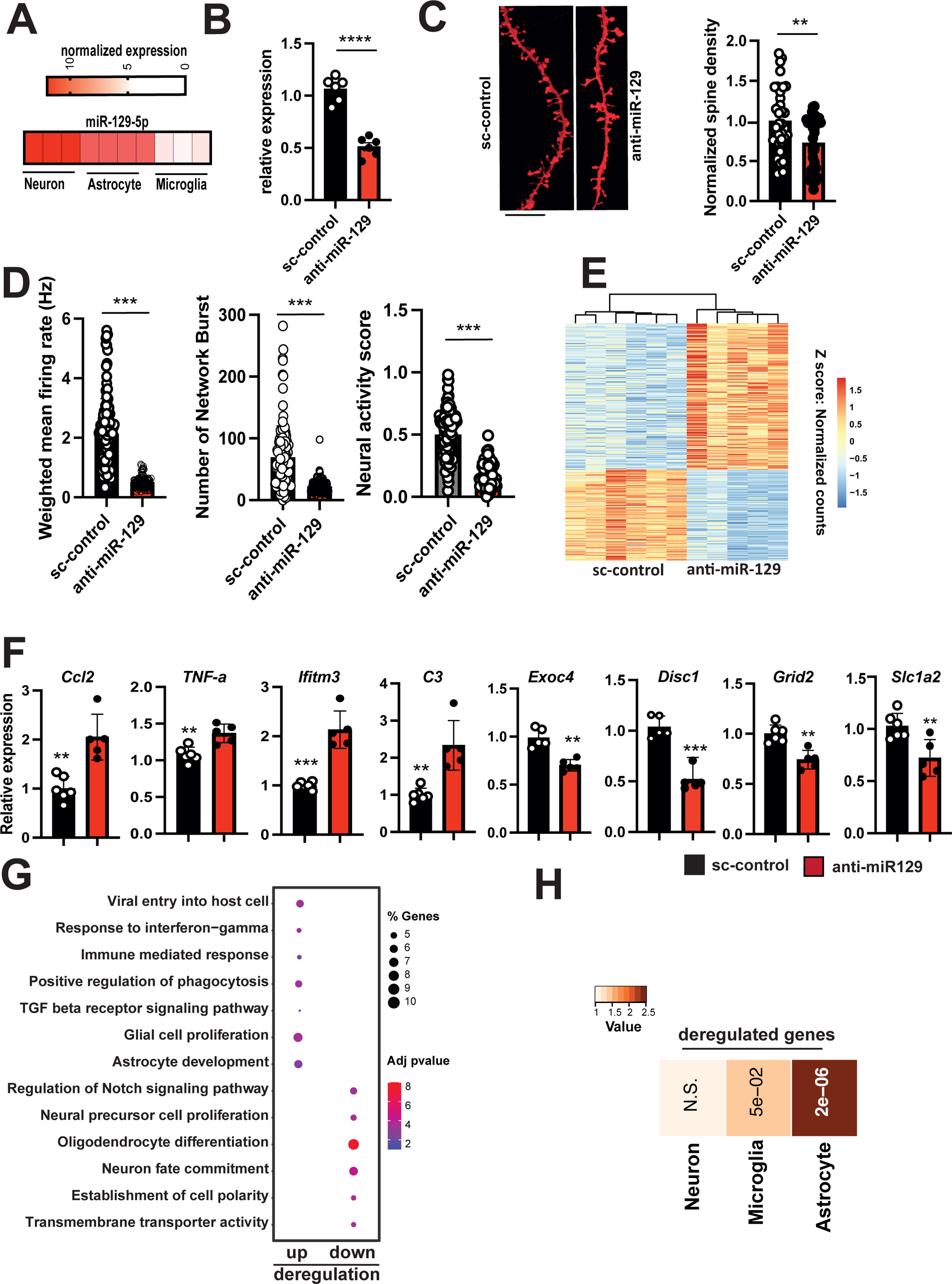
Knock down of miR-129-5p in PNM cultures impairs neuronal function and leads to deregulated gene-expression. **A.** Heat map showing the expression of miR-129-5p in different neural cell types. **B.** Bar plot showing qPCR analysis of miR-129-5p levels in PNM cultures upon anti-miR-129 administration. When compared to the control group miR-129-5p expression is significantly reduced. Data was normalized to U6 expression. \*\*\*\**P* < 0.0001, two-tailed, unpaired t-test (n=6 per group).**C**. Left panel: Representative confocal images of neuronal dendrites from sc-control and anti-miR-129 treated PNM cultures. Right panel shows the quantification of dendritic spines (number of total spines per length of a chosen dendritic segment). Anti-miR-129 treated neurons displayed a significantly reduced number of dendritic spines (number of dendritic segments analyzed: scramble control = 31, miR-129-5p inhibitor = 32. \*\**P* < 0.01, Unpaired t-test, two-tailed). **D.** Bar plots showing the results of multielectrode array (MEA) experiments in anti-miR-129-5p-treated PNM cultures compared to the sc-control group. Comparison of the weighted mean firing rate (left panel), the number of network bursts (middle panel) and the neuronal activity score (right panel) were impaired in anti-miR-129-5p-treated PNM cultures when compared to the sc-control group. A total of 29 recordings per group were conducted, with n=6 samples per group (unpaired, two-tailed t-test was performed, *P<0.05, **P<0.01, ***P<0.001). **E.** Heatmap displaying gene expression changes in PNM cultures upon anti-miR-129 treatment (log2foldchange ±0.4, and adjusted p-value <0.05). **F.** Bar plots showing qPCR results for selected up- or downregulated genes detected via RNA seq (n=6/group, two-tailed, unpaired t-test; \**P* < 0.05, \*\**P* < 0.01, \*\*\**P* < 0.001). **G.** A dot plot is presented, illustrating the GO term analysis of the differentially expressed genes shown in (E). To aid visualization, similar GO terms were clustered using GO semantic similarity, and the parental GO term was highlighted. **H**. Heat map showing the cell type enrichment of genes identified in (E) using published datasets to detect neuronal and astrocytic genes. *P*-values are based on hypergeometric testing to detect overlap between deregulated genes with published datasets from neuron, microglia and astrocytes (Fisher’s exact test, adjusted p-value with Benjamini-Hochberg (BH) correction). The color key reflects the odds ratios derived from the cell type enrichment analysis. Darker shades correspond to higher odds ratios, indicating a stronger enrichment of deregulated genes for a specific cell type, while lighter shades represent lower odds ratios, suggesting less enrichment. The visual gradation from light to dark therefore encapsulates the varying degrees of gene enrichment across different cell types. Error bars = mean ± SD.

### Reduction of miR-129-5p levels in astrocytes impair neuronal plasticity

The gene expression analysis described above suggests that reduced levels of miR-129-5p may particularly impact astrocyte function. Given the limited understanding of the role of astrocytes in FTD pathogenesis [55] we have decided to investigate the role of miR-129-5p in astrocytes in more detail.

First, we administered anti-miR-129 oligonucleotides to primary astrocyte cultures and performed RNA sequencing. Astrocytes treated with sc-control served for comparison. RNA was collected 48 hours after transfection and knock down of miR-129-5p was confirmed via qPCR **(Fig. 5A)**. Differential expression analysis revealed 704 deregulated genes, with 279 upregulated and 425 downregulated genes (basemean >= 50, log2FC +/− 0.4, adjusted p value < 0.05) (**Fig. 5B**, **Supplementary Table 10**). GO term analysis of the upregulated genes revealed processes related to neuroinflammation such as interleukin responses, while the downregulated genes were associated with processes indicative of deregulated synaptic support function, such as “Synaptic signaling”, “synapse organization” or “axon development” (**Fig. 5C**, **Supplemental Table 11)**. We confirmed the up-regulation of key cytokines Tnfa, Il-1b and Il6 via qPRC **(Fig. 5D)**. The fact that transcripts representative for GO terms linked to synaptic function are downregulated in astrocytes may appear surprising at first. However, a closer look at the affected transcripts indicate that many also have described functions in astrocytes. One example are voltage-gated calcium channels that contribute to calcium signaling in astrocytes. For example, we confirmed via qPCR the down-regulation of *Calcium voltage-gated channel subunits* (*Cacna) 1c* and *Cacna1d* that encode the alpha 1 ion-conducting pore of the Cav1.2 and Cav1.3 channels, respectively. We also confirmed the reduced expression of *Cacna2d1* and *Cacna2d3* that encode the corresponding α_2_δ subunits.

**Fig. 5.**
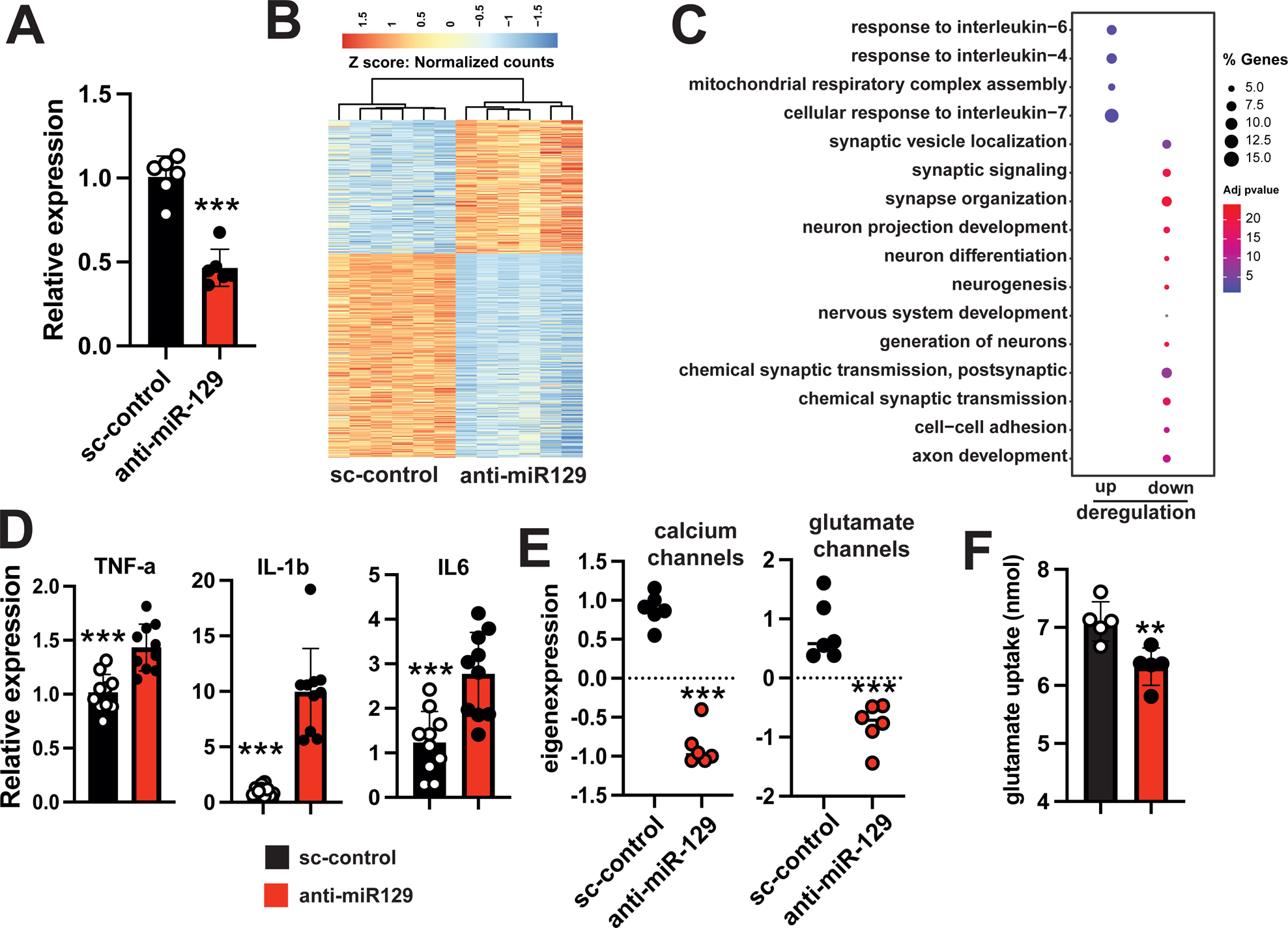
miR-129-5p controls astrocytic gene-expression, cytokine levels and glutamate uptake. **A.** qPCR analysis of miR-129-5p expression in primary astrocyte (DIV12) culture 48 h after anti-miR-129 or sc-control treatment(\*\*\**P* < 0.001 for anti-miR-129 treated cell compared to the sc-control condition, two-tailed, unpaired t-test; n=6 per group. **B**. Heatmap displaying gene expression analysis (basemean ≥ 50, log2foldchange ±0.4, and adjusted p-value <0.05) in primary astrocytes treated with anti-miR-129 or sc-control. **C**. A dot plot is utilized to illustrate the GO term analysis of the genes shown in (B). **D.** Bar plots showing qPCR analysis for inflammatory cytokines *TNF-a*, *Il1b* and *Il6*. (n=6, unpaired t-test; two-tailed, \*\*\**P* < 0.0001 for anti-miR-129 treated cells vs sc-control group). **E.**The plots display eigen-expression calculated based on qPCR results, comparing the expression of Cacna1c, Cacna1d, Cacna2d1, and Cacna2d3 (calcium channels, left panel), and Scl1a1, Slc1a2, and Scl1a3 (glutamate transporters, right panel) in anti-miR-129 treated astrocytes versus sc-control treated astrocytes (n=6, unpaired t-test; two-tailed, \*\*\**P* < 0.001). **F.** Bar plot showing the results of a glutamate uptake assay in astrocytes (n = 4, unpaired t-test; two-tailed, \*\**P* < 0.01, from anti-miR-129 treated cells vs sc-control group). Error bars indicate mean ± SD.

Since these 4 genes encode subunits of the same calcium channel, expression of these 4 genes is summarized as one eigen-value (eigen-expression) **(Fig. 5E)**. Of note the Cav1.2 and Cav1.3 channels are known to play a role in astrocytic calcium signaling, have been linked to reactive astrocyte states (PMID: 27247164) and show decreased expression upon prolonged Aß treatment (PMID: 24435206). Similarly, the *Scl1a1*, *Slc1a2* and *Scl1a3* genes that encode glutamate transporter were all downregulated in anti-miR-129 treated astrocytes. These glutamate transporters are known to be expressed in astrocytes and play a key role in synaptic glutamate homeostasis. We confirmed via qPCR the decreased expression of these glutamate transporters. The expression of *Slc1a1*, *Slc1a2* and *Slc1a3* upon miR-129-5p knock down in primary astrocytes is depicted as eigenexpression **(Figure 5E)**. In line with this observation, astrocytes treated with anti-miR-129 exhibited reduced glutamate uptake when compared to the scramble control group. **(Fig. 5F)**. These data support the hypothesis that miR-129-5p controls astrocytic processes that are known to play a role in neurodegenerative diseases including FTD.

It is important to note that changes in the levels of a single miRNA can significantly impact the cellular transcriptome. These changes occur through direct binding of the miRNA to target transcripts and subsequent secondary effects.

To gain further insight into how reduced levels of miR-129-5p may affect the transcriptional network in astrocytes, we compared the list of genes differentially expressed in astrocytes upon anti-miR-129-5p treatment with the list of experimentally confirmed miR-129-5p targets. This analysis identified 50 confirmed miR-129-5p targets **(Fig S2)**.

Using this data, we constructed a miR-129-5p transcriptional interaction network. Remarkably, 80 % of the differentially expressed genes could be explained by this network. This included genes for which down-regulation was confirmed via qPCR such as the glutamate transporter genes *Scl1a1*, *Slc1a2*, and *Scl1a3*, as well as the *Cacna1c*, *Cacna1d*, *Cacna2d1*, and *Cacna2d3* genes encoding calcium channels as well as up-regulated genes linked to neuroinflammation **(Fig S2)**.

### Astrocytic miR-129-5p controls neuronal synaptic plasticity

In summary, our data suggest that reduced levels of miR-129-5p in astrocytes increase inflammatory processes and compromise neuronal support function. Although anti-miR-129 treatment in PNM cultures impaired the number of dendritic spines and reduced neuronal network plasticity, it is likely that this effect is a combination of reduced miR-129-5p function in both neurons and astrocytes. To directly investigate the impact of reduced expression of miR-129-5p in astrocytes on neuronal plasticity, we conducted a co-culturing experiment. Primary astrocytes were treated with either a sc-control or anti-miR-129 oligonucleotides for 48 hours before being transferred to PNM cultures **(Figure 6A)**. In this experimental setup, any observed effect on neuronal plasticity is solely due to the loss of miR-129-5p in astrocytes. After 48 hours, cells were fixed, and the neuronal spine density was analyzed. Neurons grown in the presence of astrocytes in which miR-129-5p was knocked down exhibited a significantly decreased number of dendritic spines compared to PNM cultures that received sc-control treated astrocytes **(Figure 6B)**. The same experimental approach was employed to study neuronal network plasticity via MEA assays. PNM cultures were grown on MEA plates, and basal activity was assessed at DIV 10. Subsequently, the cultures were treated with astrocytes that had received anti-miR-129-5p or sc-control oligonucleotides. Neuronal activity, including parameters such as the weighted mean firing rate and the number of network bursts, was significantly impaired in PNM cultures that had received anti-miR-129-treated astrocytes **(Figure 6C)**. While these data do not exclude the possibility that loss of neuronal miR-129-5p function affects synaptic plasticity, they suggest that reduced levels of miR-129-5p in astrocytes are sufficient to cause aberrant neuronal plasticity.

**Figure 6.**
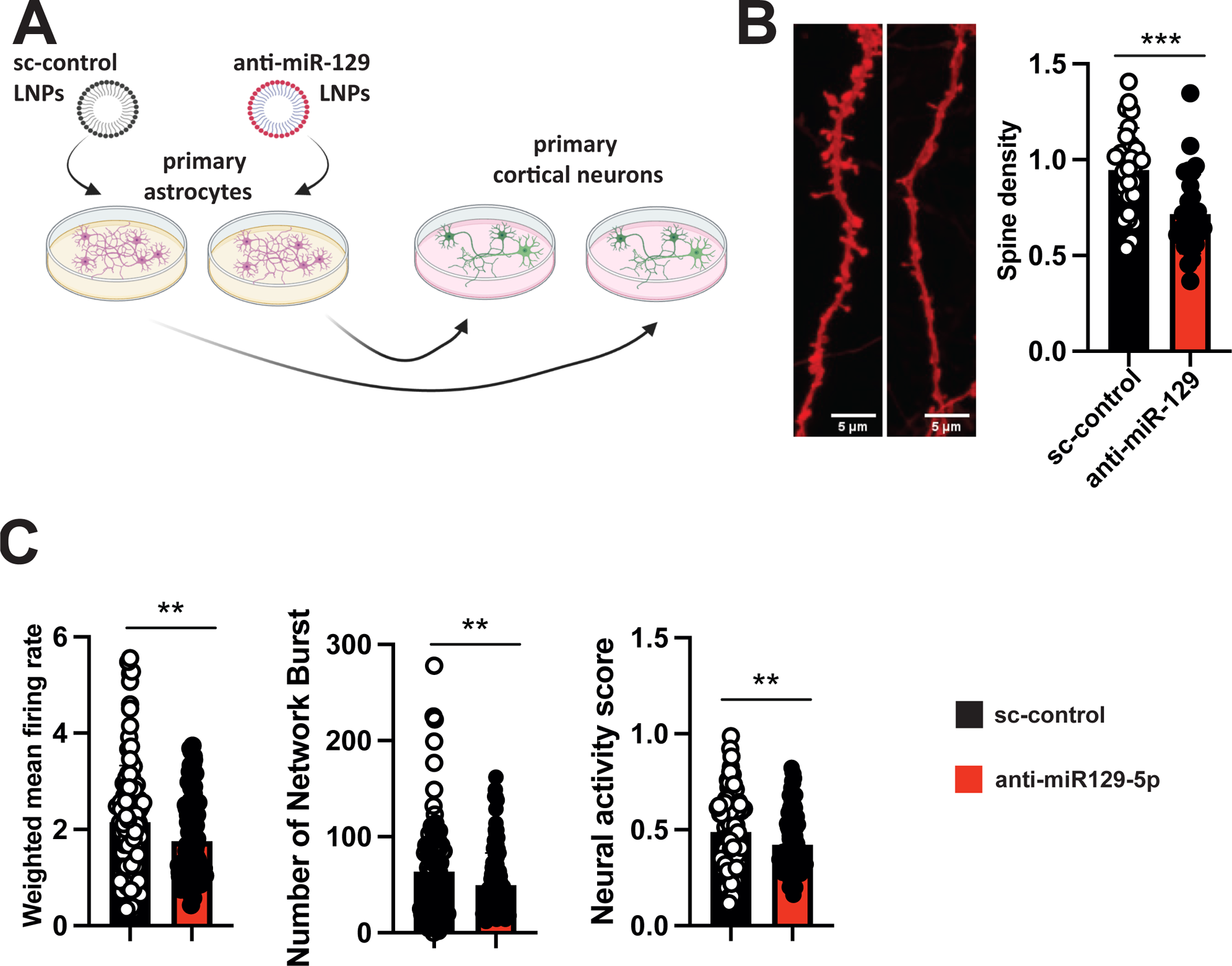
Reduced miR-129-5p levels in astrocytes affect neuronal structure and plasticity. **A.** Schematic representation of the co-culturing experiment. **B**. In the left panel, representative images of dendrites from PMN cultures are shown, to which astrocytes treated with sc-control or anti-miR-129 had been transferred. The right panel presents a bar graph quantifying dendritic spines. A total of 41 dendritic segments were analyzed for PMN cultures treated with sc-control, while 42 segments were analyzed for anti-miR-129-5p treated astrocytes (n=6/group). An unpaired, two-tailed t-test was conducted (\*\*\**P* < 0.001). **C.** PMN cultures were grown on Axion MEA plates, while primary astrocytes were initially cultured in T-75 flasks. At DIV 10, astrocytes were treated with anti-miR-129- or sc-control. At DIV 12 the astrocytes were co-cultured with the PMN cultures on the MEA plates. Spontaneous neuronal activity was recorded every hour for 10 minutes, with the entire recording session spanning 29 hours. The bar plots show the results of MEA experiments. Comparison of the weighted mean firing rate (left panel), the number of network bursts (middle panel) and the neuronal activity score (right panel) were impaired in PNM that were co-cultures with astrocytes treated with anti-miR-129. A total of 29 recordings per group were conducted, with n=6 samples per group. An unpaired, two-tailed t-test was performed (**P<0.01 anti-miR-129 vs sc-control). Error bars indicate mean ± SD.

## Discussion

In this study, we analyzed the microRNA profile in postmortem brain tissues, specifically the frontal and temporal lobes, obtained from patients with FTD who carried mutations in the *MAPT*, *GRN*, or *C9ORF72* genes. These genes are the most commonly associated with autosomal dominant inheritance of FTD. When compared to tissue from control subjects, we observed differential expression of microRNAs in all conditions. The number of deregulated microRNAs was similar in the frontal and temporal lobes of patients with MAPT mutations, as well as in the frontal lobe of patients with FTD due to C9ORF72 and GRN mutations. Relatively fewer microRNAs were deregulated in the temporal lobe of patients with FTD due to *C9ORF72* (6 microRNAs) and *GRN* (16 microRNAs) mutations. While we cannot exclude the possibility that this discrepancy is due to technical issues, it is noteworthy that previous studies have described differences in the progression of neuropathological alterations among carriers of *GRN*, *C9ORF72*, and *MAPT* mutations [32].

While data on microRNA profiling in postmortem brain tissue for FTD remains limited, our study identified several microRNAs previously linked to FTD through investigations of either cerebrospinal fluid (CSF) or postmortem brain tissue. For instance, miR-124-3p was found to be decreased in the frontal lobe of *C9ORF72* and *MAPT* carriers in our study, which is consistent with its association with synaptic dysfunction mediated by *CHMP2B* mutations, another risk gene for FTD (Gascon, 2014). Similarly, miR-204-5p was down-regulated in the frontal lobe of C9ORF72 carriers and in the temporal lobe of patients with MAPT mutations. These findings align with a previous study where decreased miR-204-5p levels were observed in CSF exosomes from presymptomatic *C9ORF72*, *MAPT*, and *GRN* carriers, suggesting that miR-204-5p could serve as a biomarker for early detection of FTD pathology [56]. Another study utilizing small RNA sequencing analyzed cortical tissue from 8 controls and 5 FTD patients, identifying miR-132-3p as down-regulated [50]. These results are consistent with another study showing decreased levels of miR-132 and miR-212 in FTD-TDP patients with or without GRN mutations [49]. MiR-132-3p, which forms a cluster with miR-212-5p, was also decreased in all analyzed tissues in our study except the frontal lobe in patients with MAPT mutations. Additionally, studies investigating CSF reported reduced levels of miR-328-3p in CSF exosomes from FTD patients [57]. We found significantly lower expression of miR-328-3p in the frontal lobe of all patients, while levels in the temporal lobe remained unaffected. Furthermore, we observed reduced miR-181c-5p levels in the frontal lobe of MAPT and C9ORF72 patients, consistent with a study on CSF exosomes reporting decreased miR-181c-5p expression in FTD patients [58].

Some microRNAs previously reported to be deregulated in FTD could not be confirmed in our study. For instance, miR-203, which was identified via the combined analysis of several mouse models for FTD and found to be up-regulated in the cortex of FTD patients with Tau pathology [27], did not show deregulation in the postmortem brain tissues analyzed in our study. Similarly, a study reported altered expression of miR-183, miR-96, and miR-182 in mice upon exposure to learning or environmental enrichment, and subsequently detected decreased levels of these microRNAs in the frontal cortex of FTD patients compared to controls [59], while we did not detect changes for these miRs. The discrepancy with our data remains unclear at present, but it is noteworthy that the post-mortem delay was much shorter in our dataset, and moreover we only focused on FTD patients with a genetic diagnosis, excluding sporadic cases.

We identified two microRNAs consistently deregulated in both analyzed brain regions across different mutation carriers, miR-212-5p and miR-129-5p, both decreased in FTD patients. As discussed earlier, miR-212-5p forms a cluster with miR-132-3p, both previously found to be decreased in the brains of FTD patients [49] [50]. Deletion of the miR-132/212 cluster impairs synaptic plasticity and memory consolidation in mice [34] [35] [36], suggesting that the decrease of miR-212-5p likely contributes to FTD pathology. Moreover, miR-212-5p was found decreased in several other neurodegenerative diseases such as AD, Huntington’s disease, and PD [37] [38] [39] [40] [41] [42] [43] [44] [45], indicating a key role in neurodegenerative diseases. Additional studies suggest that targeting miR-212-5p expression is neuroprotective in models of PD, AD, and traumatic brain injury [46] [47] [48].

In summary, these findings support the robustness of our study and confirm the deregulation of miR-212-5p in FTD patients, suggesting that therapeutic strategies targeting miR-212-5p could be a promising approach in treating neurodegenerative diseases. In addition to miR-212-5p, we consistently observed the down-regulation of miR-129-5p in the frontal and temporal lobes of patients carrying *GNR*, *C9ORF72*, or *MAPT* mutations. While the role of miR-212-5p has been relatively well-studied, information regarding miR-129-5p is limited. To our knowledge, our study is the first to report decreased miR-129-5p levels in FTD. However, reduced miR-129-5p expression was recently observed in postmortem brain samples of AD patients from the ROS/MAP cohort. Additionally, miR-129-5p expression was negatively correlated with cognitive function [51]. Another study reported decreased miR-129-5p levels in the superior temporal gyrus and entorhinal cortex when comparing controls to AD patients or PD patients [16]. These findings suggest that, similar to miR-212-5p, deregulation of miR-129-5p may play a general role in neurodegenerative processes rather than being specific to FTD. In this context it is noteworthy that especially in younger patients FTD is often misdiagnosed as psychiatric diseases such as schizophrenia, bipolar diseases or major depression [60]. As such, it is interesting that so far there is no report on the de-regulation of miR-129-5p in schizophrenia. Even in a large recent study in which 573 blood samples and 30 postmortem brains of schizophrenia patients and controls were analyzed by smallRNAseq, no de-regulation of miR-129 was observed [61]. However, it should be mentioned that recent study reported that miR-129-5p is decreased in exosomes isolated from the postmortem brain of males but not females that suffered from depression [62]

Functional data on miR-129-5p is rare. Decreased levels of miR-129-5p have been observed in various cancers and low miR-129-5p levels have been associated with a reduced efficacy of chemotherapy because miR-129-5p was found to control various genes linked to cell proliferation [63] [64]. However, little is known about the role of miR-129-5p in the brain. Therefore, we had decided to study miR-129-5p in greater detail and observed that downregulation of miR-129-5p in PMN cultures led to the up-reguation of genes linked to neuroinflammation while transcripts linked to synaptic function were decreased. Interestingly, the deregulated genes were mainly linked to astrocytes which are known to contribute to neuroinflammation and moreover orchestrate important synaptic and neuronal support functions [65]. Although most of the research in FTD has been focused on neuronal dysfunction and neuronal cell death, the role of astrocytes in the pathogenesis of FTD and other neurodegenerative diseases is increasingly being recognized [66] [67] [68] [55]. In line with these observations, decreased levels of miR-129-5p have been associated with the induction of inflammatory processes in various other diseases such as kidney injury or rheumatoid arthritis [69] [70]. Moreover, one previous study linked miR-129-5p to neuroinflammation and demonstrated that increasing the levels of miR-129-5p in a mouse model for ischemia ameliorated the expression of proinflammatory cytokines and restored motor function [71]. While neuroinflammation is mediated by multiple types of glial cells, primarily microglia, we specifically reduced miR-129-5p levels in astrocytes and in astrocytes later co-cultured with primary cortical neurons. Through this approach, we demonstrated the specific role of miR-129-5p in orchestrating astrocyte-mediated inflammation. Thus, reducing miR-129-5p levels did not only increased the expression of pro-inflammatory cytokines but also decreased the expression of transcripts linked to key neuronal support function such as glutamate-uptake, which was reduced upon miR-129-5p knock down. In line with such observations, decreasing miR-129-5p levels in astrocytes led to a reduced number of dendritic spines and reduced neuronal network activity. These findings are in agreement with previous studies showing that aberrant function of astrocytes can have detrimental effects on neuronal plasticity [72]. These data clearly show that lower miR-129-5p levels in astrocytes are sufficient to cause neuronal phenotypes observed in FTD. At the same time it is important to mention that these findings do not rule out a role for miR-129-5p in neurons or other cells of the human brain. For example neuronal miR-129-5p has been linked to epileptic plasticity and homeostatic synaptic downscaling in excitatory neurons [73].

It is therefore likely that deregulation of miR-129-5p in other cell types than astrocytes could contribute to FTD. Indeed, one limitation of our study is the fact that we analyzed bulk postmortem tissue. Since in the human brain, miR-129-5p levels are highest in neurons and astrocytes it is possible that the decreased expression in FTD could be attributed to neurons as well as astrocytes but potentially also other brain cells. It would therefore be important to study the brains from FTD patients via single cell smallRNA-seq or via semi quantitative immunohistochemical analysis in the future. Another shortcoming of our study is that the analysis has so far been limited to patients with either *C9ORF72*, *GRN* or *MAPT* mutations. Although these 3 mutations account for the majority of genetic FTD cases, it would be important to assay miR-129-5p levels in brain tissue for sporadic patients or patients with other known mutations. In this context it would also be interesting to analyze miR-129-5p expression patients suffering from ALS.

Another crucial consideration is that miRs are not only being explored as drug targets for various diseases, including brain disorders), but they are also being investigated as potential biomarkers for central nervous system (CNS) pathology when measured in cerebrospinal fluid (CSF) or blood [74] [75] [14]. Several studies have examined microRNAs in blood with the aim of identifying circulating biomarkers for FTD. While some studies focused on analyzing candidate miRs, others utilized OMICS approaches. However, to our knowledge, there have been no reports on the levels of miR-129-5p in the blood of FTD patients, despite evidence demonstrating its detectability in blood plasma, serum, or blood-derived exosomes [64]. Nonetheless, other miRs found to be dysregulated in our brain tissue analysis have been identified as potential blood-based biomarkers for FTD. For instance, decreased plasma levels of miR-127-3p and miR-502-3p were observed in FTD patients [76] [77]. Consistent with these findings, we also observed decreased miR-127-3p levels in the frontal lobe of patients with C9ORF72 and GRN mutations, as well as in the temporal lobe of patients with MAPT mutations. Similarly, miR-502-3p levels were decreased in the temporal lobe of MAPT carriers in our analysis. In a recent study, 65 miRs previously associated with FTD/ALS were investigated, and differential expression was confirmed in plasma samples from C9ORF72 or GRN carriers for 50 of these miRs (Kmetzsch, 2022). Twenty of these miRs were also detected in our study of postmortem brain tissue. Furthermore, nine miRs were found to be dysregulated in plasma when comparing controls to sporadic patients or those with GRN, C9ORF72, or MAPT mutations [78]. Among these miRs, miR-155-5p, miR-222-3p, miR-140-3p, miR-106b-3p, miR-16-5p, miR-223-3p, miR-27a, and miR-124-3p were also dysregulated in at least one of the analyzed brain tissues in our study. Finally, Magen et al. employed machine learning to identify a signature of 13 miRs measured in plasma to predict FTD. Of these miRs, nine were also dysregulated in at least one of the analyzed brain tissues from FTD patients, including miR-423, miR-125b-5p, miR-185-5p, let7d-5p, miR-107, miR-361-5p, miR-379-5p, and miR-378-5p [79].

These findings provide additional support for the robustness of our dataset, which could contribute to the establishment of miRNA signatures suitable for diagnosing FTD. Building upon these results, it would be valuable to investigate miR-129-5p levels using qPCR in blood samples obtained from both pre-symptomatic and symptomatic FTD patients carrying C9ORF72, GRN, and MAPT mutations.

## Conclusion

In summary, our findings confirm the involvement of miR-212-5p in FTD pathogenesis and suggest that targeting miR-129-5p could represent a novel therapeutic strategy for treating not only FTD but also other neurodegenerative diseases. While further research is needed to experimentally validate this hypothesis in model systems, our proposition is supported by a study investigating the effects of physical exercise in AD. Physical exercise has been shown to enhance cognitive function in both model organisms and humans, correlating with alterations in miRNA expression profiles [80]. Notably, physical exercise has been found to elevate miR-129-5p levels in the blood samples of AD patients and AD mouse models [81] [82] [83].

In conclusion our study provides a comprehensive analysis of the microRNAome in the brains of FTD patients, shedding light on potential therapeutic avenues. Additionally, we report, for the first time, a role for miR-129-5p in astrocytes and provide evidence suggesting that miR-129-5p-mediated dysregulation of astrocytic function may play a crucial role in the development of neuropathology in FTD.

## Declarations

### Ethics approval and consent to participate

RNAseq analysis of postmortem brain tissue was approved by the ethical committee of the University Medical Center Göttingen (AZ 2/8/22 and AZ 29/9/18).

### Consent for publication

All author grant consent for the publication of our manuscript

### Availability of data and material

RNAseq data from is available via GEO database. GEO accession GSE262895. smallRNA-sequencing data from the human tissue samples will be made available via the European Genome-phenome Archive(EGA).

### Authors’ contributions

LK performed experiments, analyzed and interpreted RNA-seq data, designed all experiments, and drafted the paper. RP performed MEA assay and performed some qPCR analysis. SS performed glutamate uptake assay and provided support in astrocyte culture. SB and ALS provided technical support and performed RNAseq. ALS performed RNA isolation. DMK and TP provided bioinformatics support. PH provided FTD samples. FS and AF arranged funding, designed and supervised the study, drafted, and revised the final version of the manuscript.

## Acknowledgements

n/a

---

**Fig. S1.**
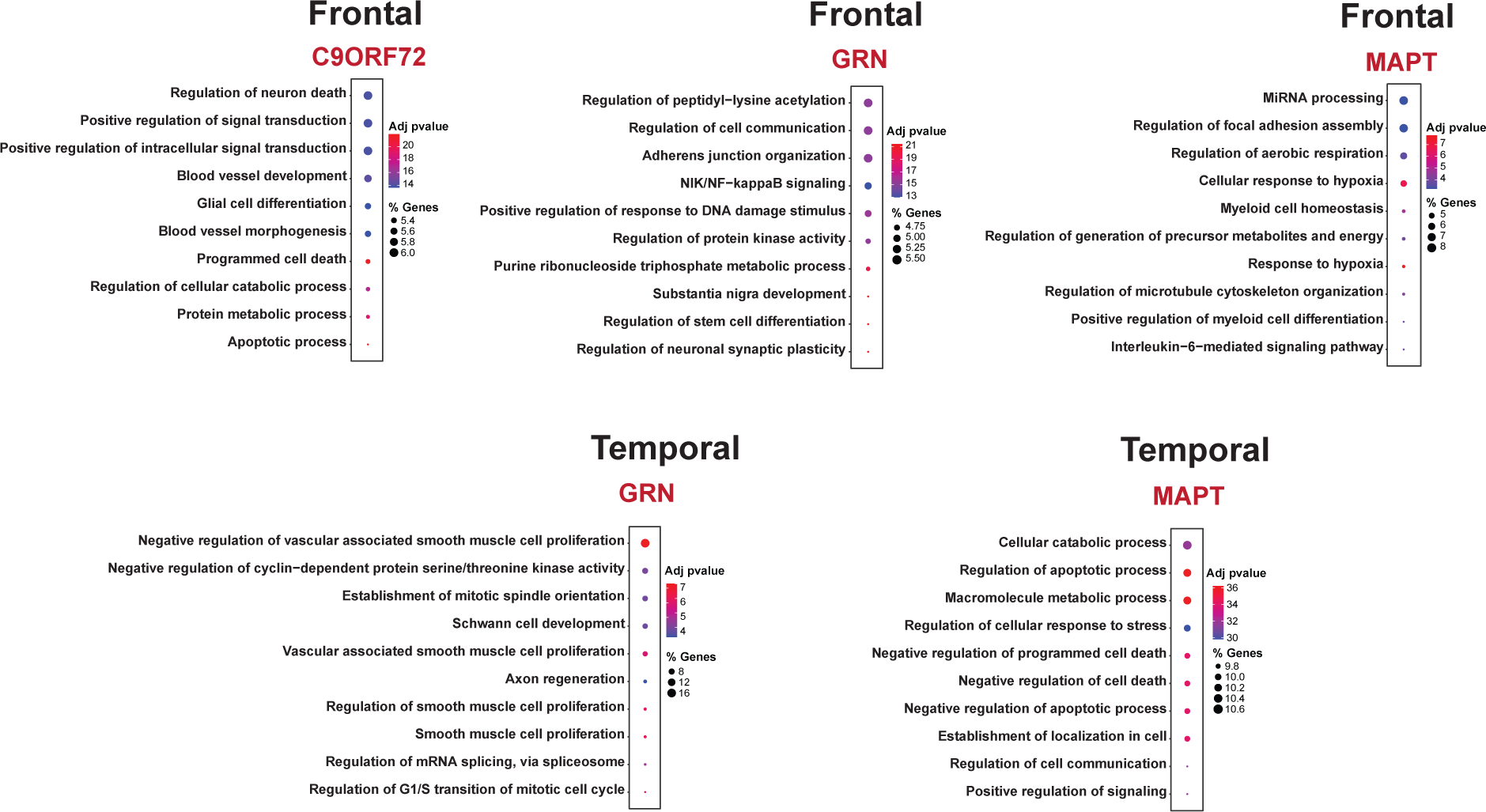
Pathways Affected by miRNAs Specifically Deregulated in Response to either *C9ORF72*, *GNR* or *MAPT* mutations and the frontal or temporal lobe. The graphs display the top 10 enriched GO terms (Biological processes) identified based on brain-expressed target transcripts of miRNAs deregulated exclusively in the depicted condition. It’s important to note that no GO term analysis was conducted for the temporal lobe in the case of *C9ORF72* carriers, as only one miRNA was deregulated in this condition

**Fig. S2.**
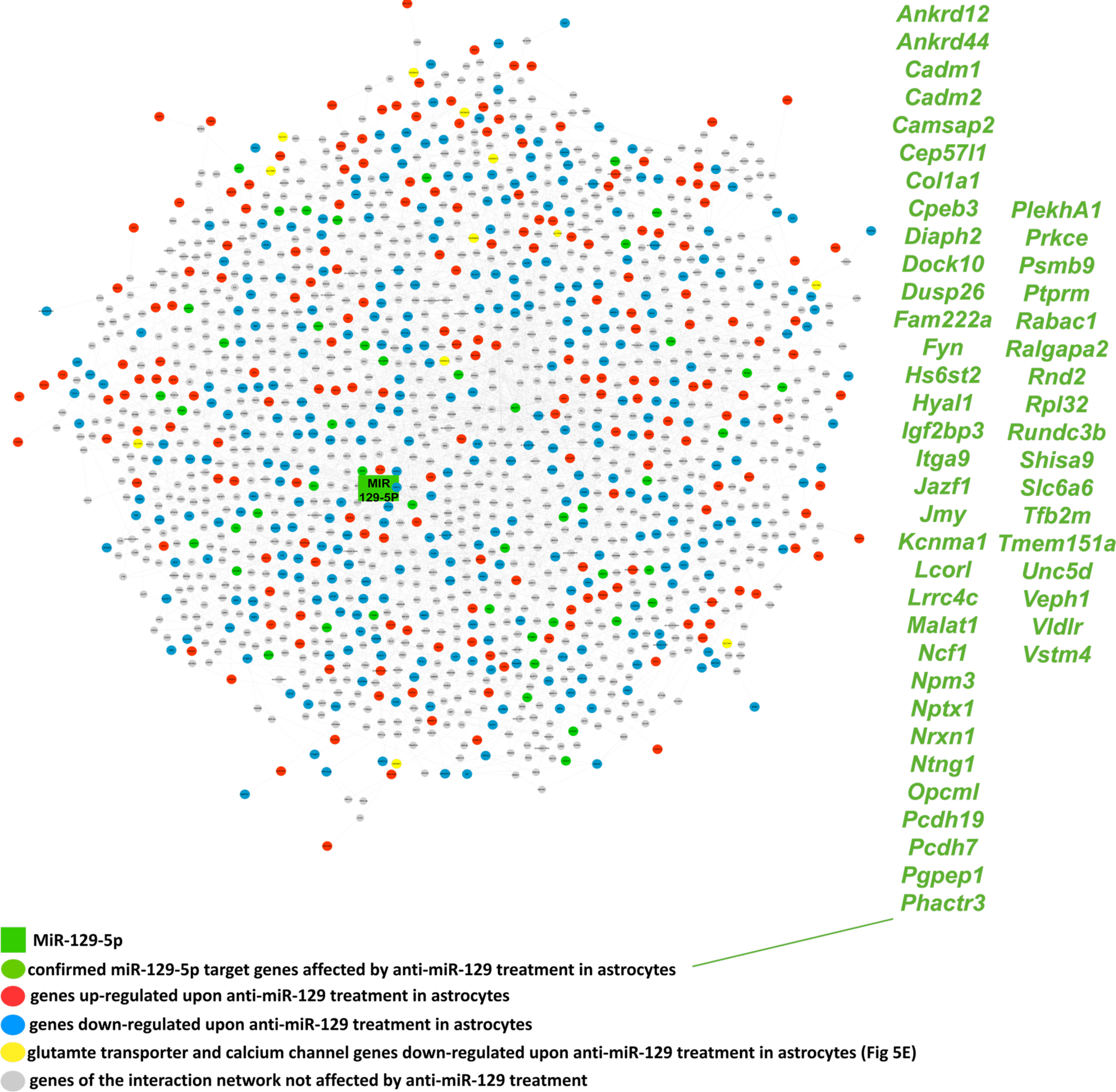
mIR-129-5p interaction network. We identified 50 confirmed miR-129-5p target genes (shown in green) among the transcripts deregulated in astrocytes upon miR-129-5p knockdown. Using these data, we built a gene expression interaction network that could explain approximately 80% of the transcripts detected as differentially expressed in the corresponding RNA-seq data. In yellow, we highlight the genes that were downregulated upon miR-129-5p knockdown and encode glutamate transporters and calcium channels, confirmed via qPCR (see Fig. 5E).

